# Laterality of subcortical structures predicts spontaneous brain dynamics

**DOI:** 10.64898/2026.07.13.738145

**Authors:** Tara Ghafari, Andrew J Quinn, Ole Jensen

**Affiliations:** Department of Experimental Psychology, University of Oxford, UK; Centre for Integrative Neuroscience, Oxford Centre for Human Brain Activity, Department of Psychiatry, University of Oxford, UK; Centre for Human Brain Health, School of Psychology, University of Birmingham, Birmingham, UK

## Abstract

Subcortical structures contribute to neocortical activity through extensive circuits supporting cognition and behaviour. Although these anatomical connections are well established, how subcortical structure relates to neocortical oscillations remains poorly understood. Previous studies have linked subcortical volume asymmetries to cortical alpha modulation during attention tasks, but whether these relationships extend to spontaneous brain activity remains unclear. Here, we investigated whether hemispheric asymmetries in subcortical volume are associated with lateralised resting-state oscillatory power. We analysed structural MRI and resting-state magnetoencephalography (MEG) data from 532 healthy adults in the Cambridge Centre for Ageing and Neuroscience (CamCAN) cohort, correlating volume asymmetries of seven subcortical structures with MEG power asymmetries across four frequency bands. We found that globus pallidus asymmetry was positively associated with alpha-band power lateralisation, extending previous task-based findings to the resting state. Within the striatum, putamen asymmetry was positively associated with frontal beta-band lateralisation, whereas caudate asymmetry was negatively associated with posterior beta- and delta-band lateralisation. Hippocampal asymmetry was negatively associated with frontal delta-band lateralisation. Despite the complexity of cortico-subcortical connectivity, these findings reveal distinct frequency-specific relationships between subcortical anatomy and spontaneous neocortical activity that align with the established functional roles of these structures. Our results demonstrate that these relationships can be detected non-invasively in resting state without requiring direct recording of deep brain regions, providing a foundation for investigating whether resting-state MEG can identify early functional changes associated with subcortical neurodegeneration.

## Introduction

Subcortical structures are anatomically positioned to shape neocortical computation. Through direct and indirect cortico-subcortical loops, the basal ganglia, thalamus, and hippocampus contribute to the organisation of cortical activity supporting attention, action selection, memory, and cognitive control (Alexander et al., 1986; Buzsáki, 2006; Haber, 2003). Although these pathways are well described anatomically, it remains less clear how individual differences in subcortical structure are expressed functionally in neocortical activity, and whether such interactions can be detected non-invasively with magnetoencephalography (MEG).

A growing body of work suggests that hemispheric asymmetry provides a useful framework for probing these structure–function relationships. In a reward-related spatial attention task, hemispheric asymmetry of the volume of the globus pallidus (GP) detected using MRI predicted hemispheric differences in posterior alpha modulation on MEG. This suggests that the GP contributes to the control of neocortical alpha activity (Mazzetti et al., 2019). Extending this approach, it was demonstrated that asymmetries of the GP, caudate nucleus (CN), and thalamus predicted attention-related alpha power modulation (Ghafari et al., 2024). More recently, hemispheric asymmetry of the putamen was shown to predict individual differences in pseudoneglect, linking striatal structure to spatial attentional bias in healthy adults (Ghafari et al., 2025). Together, these findings suggest that naturally occurring subcortical asymmetries are related to lateralised neocortical computations.

The present study asks whether this principle generalises beyond task-evoked attention to resting-state oscillatory activity. This is important because resting-state MEG offers a clinically attractive route to probing cortico-subcortical systems without requiring task performance, which is particularly relevant for aging and neurodegenerative populations. Neurodegenerative disorders such as Alzheimer’s disease, Parkinson’s disease, and mild cognitive impairment involve progressive changes in subcortical structures, including the hippocampus and basal ganglia (Braak & Braak, 1991; Jack Jr et al., 1997; Li et al., 2022; Zeighami et al., 2015).

However, structural MRI typically detects atrophy only after substantial tissue loss has occurred (Frisoni et al., 2010; Heim et al., 2017). Functional measures such as EEG and MEG may therefore provide earlier evidence of altered neocortical neural communication, than structural changes in subcortical structures, although MEG is relatively insensitive to deep sources because subcortical signals are attenuated and distorted before reaching the sensors (Hämäläinen, 1993; Krishnaswamy et al., 2017).

Rather than attempting to measure subcortical activity directly, we asked whether subcortical structure is reflected in lateralised neocortical oscillatory power. We used resting-state MEG data which is particularly advantageous in clinical settings where recording time is limited and patients may not be able to complete more demanding tasks. We focused on hemispheric asymmetry for two reasons. First, asymmetry is biologically meaningful: many subcortical systems show lateralised organisation, and neurodegenerative disorders often affect subcortical regions asymmetrically (Kaasinen, 2016; Minkova et al., 2017; Wachinger et al., 2016). Second, asymmetry provides a within-participant measure that reduces confounds related to head size, global brain volume, and overall signal amplitude. We hypothesised that if subcortical structures contribute to organising neocortical rhythms, then individual differences in lateralised subcortical volume should predict individual differences in lateralised MEG power.

In this study, we analysed resting-state MEG and structural MRI from the CamCAN cohort to test this hypothesis in a large sample of healthy adults. Building on previous evidence linking the GP and CN to alpha modulation, and the putamen to spatial attentional bias, we examined whether lateralised volumes of seven subcortical structures are associated with lateralised oscillatory power across canonical frequency bands. We expected the most consistent effect to involve the GP and alpha-band activity, reflecting converging evidence from previous task-based MEG studies. We further tested whether striatal and hippocampal asymmetry also relate to oscillatory activity. Establishing these relationships in healthy participants provides a foundation for future work using MEG-derived asymmetry measures as functional markers of subcortical change in aging and neurodegenerative disease.

## Method and Materials

### Participants and Dataset

Data were obtained from the Cambridge Centre for Ageing and Neuroscience (CamCAN) Stage-II release, an open-access neuroimaging dataset comprising healthy adults aged 18–88 years (Shafto et al., 2014; Taylor et al., 2017). This release includes high-resolution T1-weighted anatomical MRI and resting-state MEG recordings. A total of 590 participants were initially included. After quality control steps for both MRI and MEG, 532 participants were retained for the final analysis.

### Structural MRI: Subcortical Segmentation and Lateralisation

T1-weighted structural scans (1 mm³ MPRAGE) were processed using the FMRIB Software Library (FSL, version 6.0.7). Automated segmentation of subcortical structures was performed using the FSL-FIRST tool, which employs a Bayesian appearance model to delineate bilateral subcortical structures. The seven regions included were: caudate, putamen, globus pallidus, thalamus, hippocampus, amygdala, and nucleus accumbens.

Volumes for each structure were extracted using a custom parser. Outlier detection removed participants in whom the volume of at least one side of one subcortical structure fell below the 10th percentile, because very small volumes may reflect segmentation failure or poor anatomical fit. This step excluded 24 participants, leaving 566 for volumetric analysis. and a lateralisation Volume (LV) was computed for each structure as:

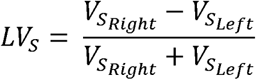

Where *V_SRight_* and *V_SLeft_* denote the voxel-based volume of a subcortical regions in the right and left hemispheres, respectively. Lateralisation indices were stored and used in all downstream analyses.

To examine the distribution and asymmetry of subcortical volumes, we plotted histograms of the lateralisation indices for all subcortical structures (Figure 1).

**Figure 1.**
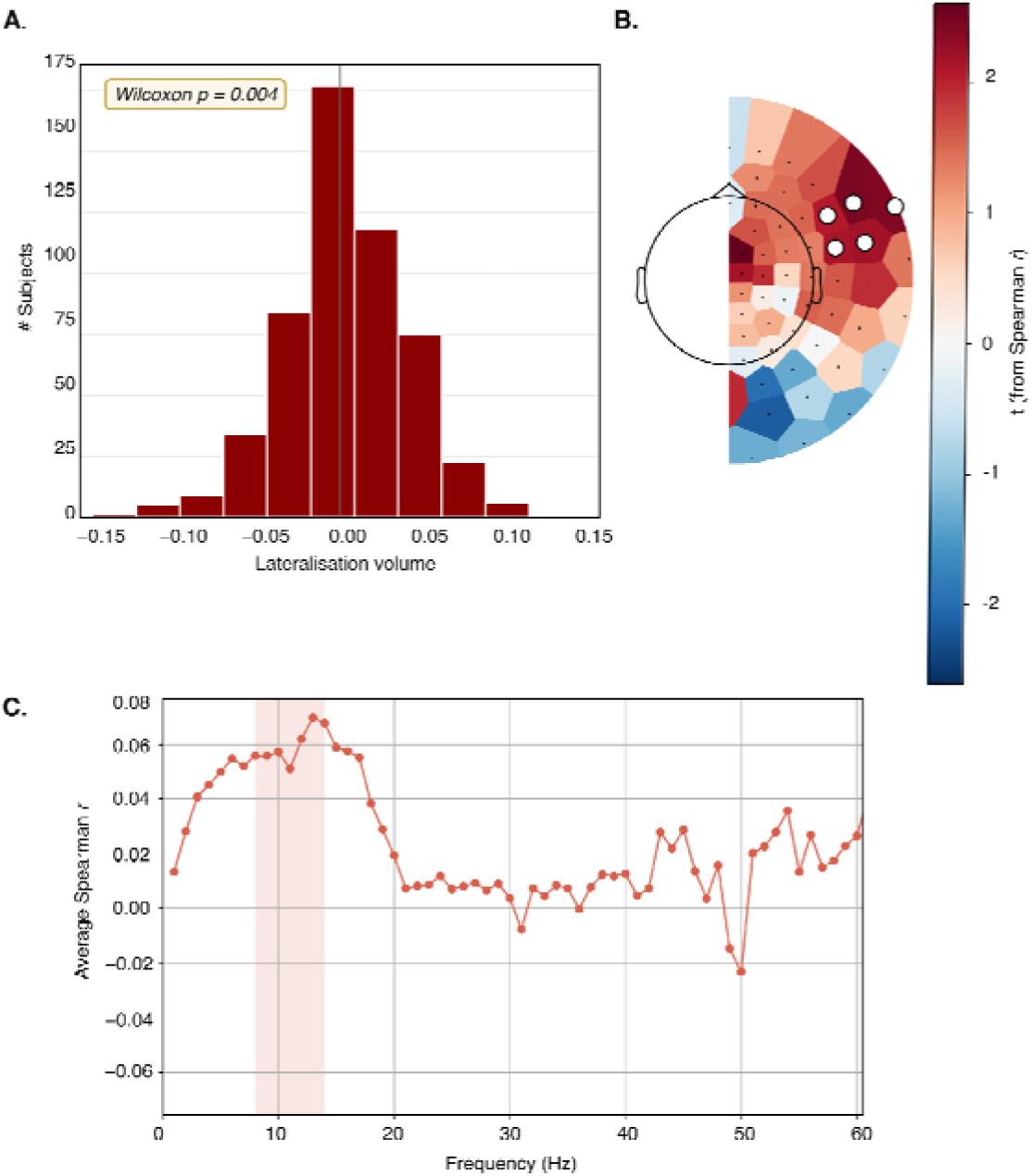
Globus pallidus lateralisation and alpha-band power asymmetry. A) Distribution of globus pallidus lateralised volume across participants (N = 532). Positive values indicate greater right than left volume, whereas negative values indicate greater left than right volume. The globus pallidus showed significant leftward lateralisation (Wilcoxon rank sum p = 0.004). B) Topographic map showing the correlation between globus pallidus lateralised volume and alpha-band power lateralisation over temporal sensors measured with combined planar gradiometers. Colours show t-transformed Spearman correlation values. White dots mark sensors within clusters that remained significant after permutation-based correction for multiple comparisons across sensors and frequency bands. C) Frequency-resolved Spearman correlation values between globus pallidus lateralised volume and the average power lateralisation across the significant sensors shown in B. The highlighted section indicates the frequency range surviving cluster correction. Together, these results show that pallidal asymmetry is selectively associated with alpha-band power lateralisation over temporal sensors.

### MEG Preprocessing and Power Lateralisation Index

We analysed eyes-closed resting-state recordings acquired with a 306-channel Elekta Neuromag VectorView system (102 magnetometers, 204 planar gradiometers). The dataset provided MaxFiltered and movement-compensated recordings, ensuring data quality prior to analysis. For each participant, a single fixed-length epoch of 7 minutes and 50 seconds was extracted.

Power spectral density (PSD) was estimated for each sensor using Welch’s method segmenting the data in 2-second windows, with 50% overlap between consecutive segments (1-second overlap). A Hamming window was applied to each segment prior to Fourier transformation. The resulting power spectra were averaged across segments, yielding a frequency resolution of 0.5 Hz.

Left–right homologous sensor pairs were defined according to the Elekta layout. For each frequency bin, a power lateralisation index (PLI) was computed as:

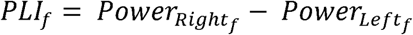

Here *Power_Right f_* and *Power _Left f_* refer to the spectral power at frequency f for the right and left sensors, respectively. For frequency band analyses, PSD estimates were averaged into canonical frequency ranges: delta (1–4 Hz), theta (4–8 Hz), alpha (8–14 Hz), and beta (14–40 Hz). PLI values were then derived for each band. Prior to computing the PLI, planar gradiometers were combined by averaging their power. Note that we did not normalise the lateralisation measure with the sum of the power as this would put more weight to higher frequencies as the power decrease with frequency.

At this stage, 34 participants were excluded due to data corruption or poor signal quality in preprocessing, leaving a final MEG–MRI-valid cohort of 532 participants.

### Volume-Power Correlation

Volume–power correlations were assessed in the 532 participants with valid MRI and MEG data. For each sensor pair and frequency band, Spearman’s rank correlation was computed between the power lateralisation index (PLI) and the lateralisation volume (LV) of each of the seven subcortical structures across participants.

These calculations yielded subcortical volume–sensor power correlation maps, producing a total of 7 (subcortical structures) × 2 (sensor types: gradiometers and magnetometers) × 4 (frequency bands) topographic representations.

### Cluster Permutation Testing

To correct for multiple comparisons across MEG sensor-pairs and frequency bands, we performed a non-parametric cluster-based permutation analysis on the sensor- wise Spearman correlations between LVs and PLIs. Correlation coefficients were converted to *t*-statistics and thresholded using a two-tailed significance level (α = 0.05). For each frequency band, neighbouring significant sensors were grouped into spatial clusters using the MEG sensor adjacency matrix, and each cluster was assigned a cluster statistic equal to the sum of its *t*-values.

Cluster significance was assessed using 1,000 permutations, in which the pairing between participants’ LVs and PLIs was randomly shuffled. For each permutation, the largest positive and largest negative cluster statistics were retained to generate null distributions controlling the family-wise error rate. Observed clusters were considered significant if their cluster statistic exceeded the 95th percentile of the positive null distribution or fell below the 5th percentile of the negative null distribution. Only sensors belonging to significant clusters were included in subsequent visualisation and spectral analyses. Full implementation details are provided in the Supplementary Methods.

## Results

We tested whether hemispheric asymmetries in subcortical volume were related to hemispheric asymmetries in resting-state MEG power. For each participant, lateralised volume (LV) was calculated for each subcortical structure as the difference between right and left volume. Similarly, power lateralisation index (PLI) was calculated for homologous left–right MEG sensor pairs across four frequency bands: delta, theta, alpha, and beta. In the main Results, we focus on combined planar gradiometers because they have a more focal sensitivity profile and are preferentially sensitive to neocortical sources close to the sensor array, whereas magnetometers are more sensitive to distant and spatially distributed sources (Garcés et al., 2017). We report the magnetometers in the supplementary materials (Supplementary figure 1). For each subcortical structure and frequency band, we correlated LV with PLI across participants and used cluster permutation testing to identify spatially contiguous significant sensor clusters.

### Globus pallidus asymmetry is associated with alpha-band power lateralisation

The globus pallidus was significantly left-lateralised in volume across participants (Figure 1A; Wilcoxon rank sum p = 0.004). We next asked whether this pallidal structural asymmetry was associated with lateralised MEG power. After cluster permutation, a significant (p-value= 0.028; controlling for multiple comparison over sensors and frequency bands) positive correlation was observed between globus pallidus LV and alpha-band PLI over temporal gradiometer sensors (Figure 1B). This indicates that participants with stronger lateralisation of pallidal volume also showed stronger lateralisation of alpha-band power in the same direction. The frequency- resolved correlation analysis confirmed that the effect was centred in the alpha range, with the significant frequencies highlighted in Figure 1C.

### Striatal asymmetries are associated with beta-band power lateralisation

We next examined whether lateralised volumes of the striatum were associated with beta-band power lateralisation. The putamen was significantly left-lateralised in volume across participants (Figure 2A; Wilcoxon rank sum p = 0.011). Putamen LV was positively correlated with beta-band PLI (p-value = 0.035) in a cluster of frontal gradiometer sensors (Figure 2B). The frequency-resolved analysis showed that this positive association was broad but centred within the beta range (Figure 2C). No significant putamen correlations were found in the delta, theta, or alpha bands.

**Figure 2.**
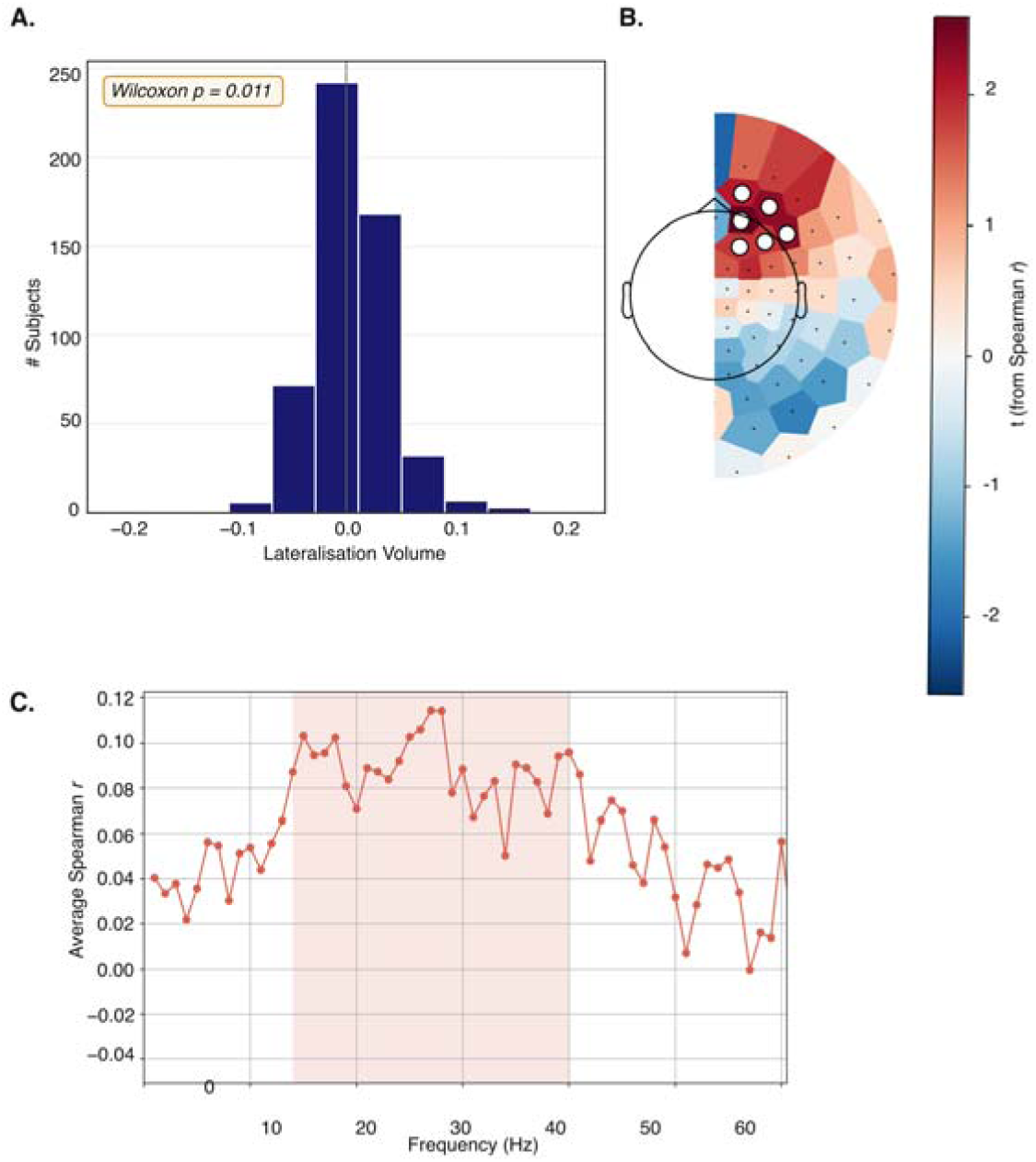
Putamen lateralisation and beta-band power asymmetry. A) Distribution of putamen lateralised volume across participants (N = 532). Positive values indicate greater right than left volume, whereas negative values indicate greater left than right volume. The putamen showed significant leftward lateralisation (Wilcoxon rank sum p = 0.011). B) Topographic map showing the correlation between putamen lateralised volume and beta-band power lateralisation measured with combined planar gradiometers. Colours show t-transformed Spearman correlation values. White dots mark sensors within clusters that remained significant after permutation-based correction for multiple comparisons across sensors and frequency bands. C) Frequency-resolved Spearman correlation values between putamen lateralised volume and the average power lateralisation across the significant sensors shown in B. The highlighted section indicates the frequency range surviving cluster correction. Together, these results show a positive association between putamen asymmetry and beta-band power lateralisation over anterior sensors.

The caudate nucleus was significantly right-lateralised in volume (Figure 3A; Wilcoxon rank sum p < 0.001). Caudate LV showed a significant negative correlation with beta-band PLI (p-value = 0.011) and with delta band PLI (p-value = 0.044) over posterior gradiometer sensors (Figure 3B, right). The frequency-resolved analysis showed that this effect was present across the beta range (Figure 3C).

**Figure 3.**
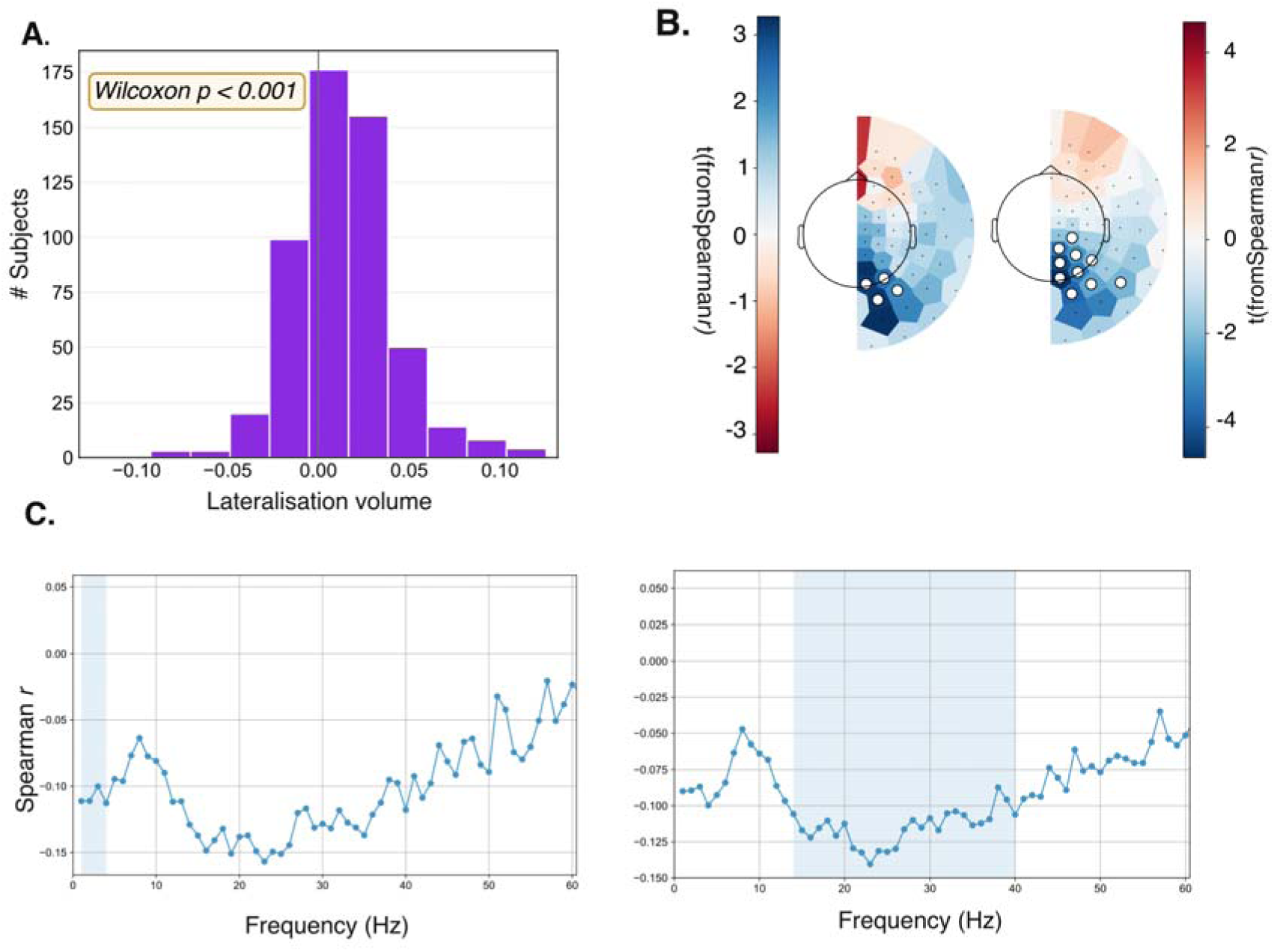
Caudate lateralisation and delta- and beta-band power asymmetry. A) Distribution of caudate lateralised volume across participants (N = 532). Positive values indicate greater right than left volume, whereas negative values indicate greater left than right volume. The caudate showed significant rightward lateralisation (Wilcoxon rank sum p < 0.001). B) Topographic maps showing correlations between caudate lateralised volume and power lateralisation measured with combined planar gradiometers. The left map shows the delta-band effect, and the right map shows the beta-band effect. Colours show t-transformed Spearman correlation values White dots mark sensors within clusters that remained significant after permutation-based correction for multiple comparisons across sensors and frequency bands. C) Frequency-resolved Spearman correlation values between caudate lateralised volume and the average power lateralisation across the significant sensors shown in B. Highlighted sections indicate frequency ranges surviving cluster correction. Together, these results show that caudate asymmetry is negatively associated with both delta- and beta-band power lateralisation over posterior sensors.

### Hippocampal asymmetry is associated with delta -band power lateralisation

The hippocampus was significantly right-lateralised in volume across participants (Figure 4A; Wilcoxon rank sum p < 0.001). Hippocampal LV was negatively correlated with delta-band PLI (p-value = 0.026) in a frontal cluster of gradiometer sensors (Figure 4B). The frequency-resolved analysis confirmed that this negative association was specific to the delta range (Figure 4C). No significant hippocampal correlations were observed in the theta, alpha, or beta bands in the gradiometer analysis.

**Figure 4.**
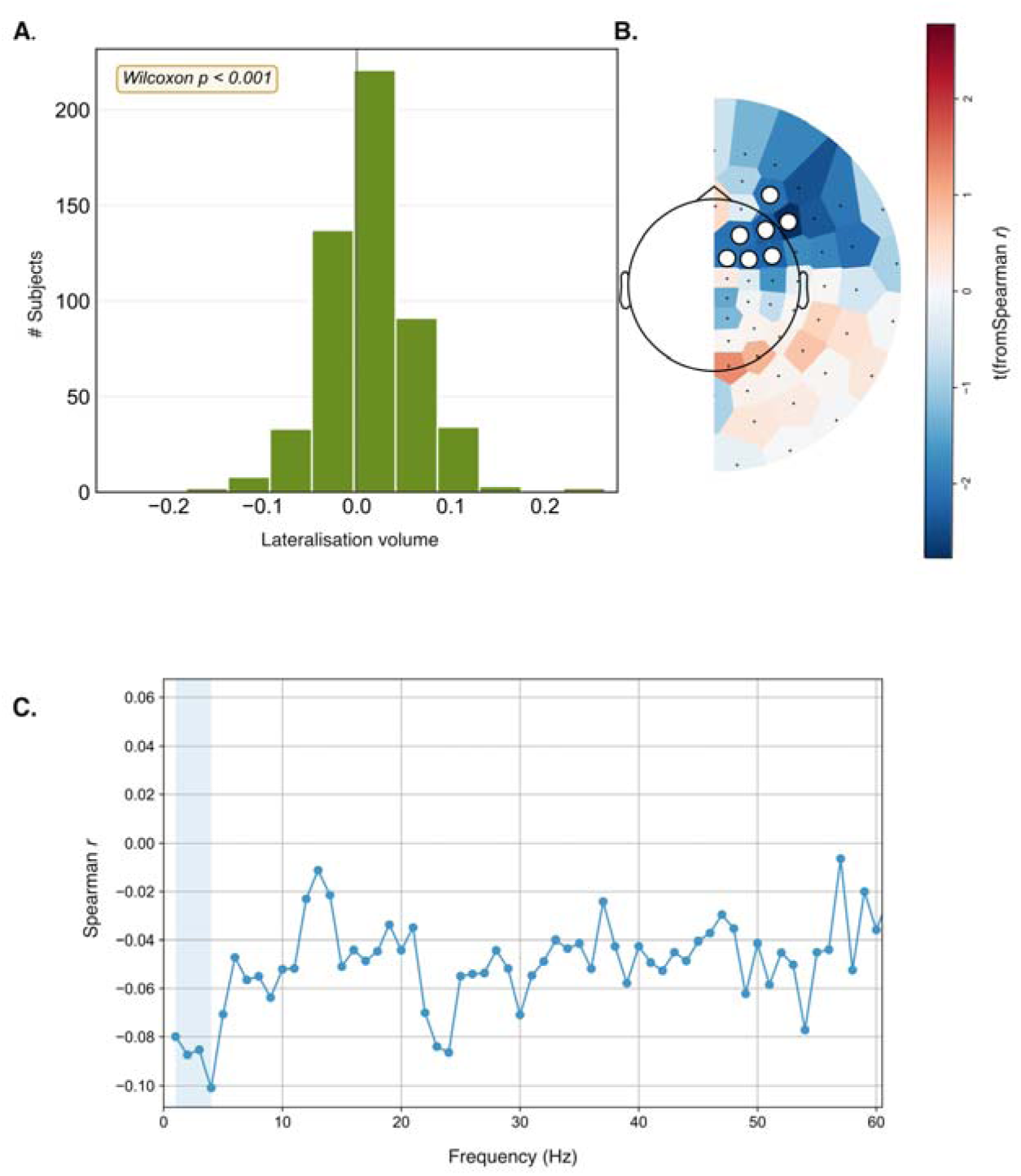
Hippocampal lateralisation and delta-band power asymmetry. A) Distribution of hippocampal lateralised volume across participants (N = 532). Positive values indicate greater right than left volume, whereas negative values indicate greater left than right volume. The hippocampus showed significant rightward lateralisation (Wilcoxon rank sum p < 0.001). B) Topographic map showing the correlation between hippocampal lateralised volume and delta-band power lateralisation measured with combined planar gradiometers. Colours show t- transformed Spearman correlation values. White dots mark sensors within clusters that remained significant after permutation-based correction for multiple comparisons across sensors and frequency bands. C) Frequency-resolved Spearman correlation values between hippocampal lateralised volume and the average power lateralisation across the significant sensors shown in B. The highlighted section indicates the frequency range surviving cluster correction. Together, these results show a negative association between hippocampal asymmetry and delta-band power lateralisation over anterior sensors.

## Discussion

We have here related hemispheric asymmetries in the volume of cortical structures to hemispheric asymmetries (lateralisation) in resting state activity as measured by MEG. We found that globus pallidus asymmetry was associated with alpha-band lateralisation, putamen and caudate asymmetries were associated with beta-band lateralisation, and hippocampal and caudate asymmetries were associated with delta-band lateralisation. These findings demonstrate that hemispheric variation in subcortical anatomy is reflected in resting-state neocortical oscillatory power in specific frequency bands, extending previous evidence for cortico-subcortical interactions beyond task-evoked activity.

### Globus pallidus asymmetry and alpha-band activity

Previous work has linked subcortical asymmetry to lateralised cortical function. Hemispheric asymmetry of the globus pallidus predicts alpha modulation during reward-guided attention (Mazzetti et al., 2019), and we subsequently found that asymmetries of the globus pallidus, caudate and thalamus predict alpha modulation during the allocation of visuospatial attention (Ghafari et al., 2024). The present findings extend these observations by showing that the relationship between globus pallidus asymmetry and cortical alpha activity is also evident at rest.

Alpha oscillations represent the dominant rhythm supporting functional inhibition and attentional gating, regulating the excitability of cortical populations according to behavioural demands (Jensen & Mazaheri, 2010; Klimesch et al., 2007). The globus pallidus is increasingly recognised as an integrative node within basal-ganglia circuits rather than simply a motor output structure (Yoshida & Hikosaka, 2025). The globus pallidus contributes to reward processing (Fiore et al., 2018; Hong & Hikosaka, 2008; Münte et al., 2017) and integrates information within cortico-basal- ganglia-thalamic loops (Giossi et al., 2024). The present findings suggest that pallidal anatomy is related not only to task-evoked alpha modulation but also to the spontaneous organisation of neocortical alpha activity.

This relationship may also have relevance for disorders characterised by attentional dysfunction. Attention-deficit/hyperactivity disorder (ADHD) has been associated with altered alpha modulation, including attenuated task-related alpha modulation and abnormal alpha lateralisation (Lenartowicz et al., 2018; ter Huurne et al., 2013), as well as structural abnormalities of the basal ganglia (Frodl & Skokauskas, 2012). The present globus pallidus–alpha relationship therefore provides a normative framework for testing whether alterations in pallidal anatomy are accompanied by changes in cortical alpha organisation and attention in individuals diagnosed with ADHD.

### Striatal asymmetry and beta-band activity

In contrast, the association between striatal asymmetry and beta-band lateralisation is consistent with extensive evidence identifying beta oscillations as a prominent feature of cortico-basal ganglia communication. Beta activity is expressed throughout basal ganglia–cortical circuits (Brown, 2007; Mallet et al., 2008). Interestingly, the putamen and caudate showed opposite correlation directions and distinct sensor topographies. This distinction may relate to their partially dissociable anatomical connectivity and functional roles within corticostriatal circuits (Alexander et al., 1986; Brovelli et al., 2011). Our findings extend this literature by showing that beta lateralisation also covaries with normal anatomical asymmetry of the striatum.

This relationship is particularly relevant to Parkinson’s disease, in which degeneration of basal ganglia circuits is accompanied by abnormal beta synchronisation. Beta-band measures are increasingly investigated as physiological markers of disease state and treatment response, with beta activity showing associations with motor symptom severity and treatment-related improvement. In particular, high-beta activity has shown promise in predicting clinical responses to therapy and informing beta-guided and adaptive deep brain stimulation (Morelli & Summers, 2023; Petersson et al., 2020). The present findings raise the possibility that cortical beta asymmetry may provide a marker of early disease related changes or degeneration of the striatal circuit, although this will require direct testing in patients.

### Hippocampal and caudate asymmetry and delta-band activity

Hippocampal asymmetry was associated selectively with delta-band lateralisation. Human intracranial recordings suggest that memory related theta oscillations observed in rodents occur at slower frequencies in the 2-4 Hz band. For instance, human iEEG studies have identified hippocampal oscillations around 3-4 Hz reflecting memory processing (Jacobs, 2014; Lega et al., 2012; Rudoler et al., 2023;

Watrous et al., 2011). Thus, the association observed here in the 1-4 Hz band is compatible with the frequency range of human hippocampal memory-related oscillations. The frontal distribution of this effect is consistent with a neocortical contribution potentially mediated by hippocampal–prefrontal interactions (Gordon, 2011)

This association may be particularly relevant to ageing and Alzheimer’s disease, where hippocampal degeneration is an early structural feature and resting-state EEG/MEG commonly shows increased slow-frequency activity. Delta and theta abnormalities are among the reproducible electrophysiological changes reported in MCI and Alzheimer’s disease (Babiloni et al., 2021; Nakamura et al., 2018).

Establishing the hippocampus–delta relationship in healthy adults therefore provides a normative basis for testing whether cortical slow-wave asymmetry tracks hippocampal degeneration in MCI and Alzheimer’s disease.

The delta-band association was not restricted to the hippocampus, as caudate asymmetry was also associated with posterior delta lateralisation. This suggests that delta-band cortical asymmetry may reflect structural variation across more than one subcortical system.

### Limitations

The present findings also highlight opportunities for further investigation. Given the broad age range of the CamCAN cohort, future analyses could examine how age- related changes in subcortical anatomy and oscillatory activity contribute to these relationships. Furthermore, longitudinal and clinical studies will be important to establish whether these associations track structural changes over time and whether they can serve as sensitive markers of early neurodegeneration.

## Conclusion

In summary, hemispheric asymmetries of subcortical anatomy were associated with frequency-specific asymmetries of resting-state cortical oscillations, with globus pallidus asymmetry related to alpha activity, striatal asymmetry predominantly to beta activity, and hippocampal asymmetry to delta activity. These findings extend previous task-based observations by showing that relationships between subcortical anatomy and cortical oscillations are also evident during spontaneous activity. Resting-state oscillatory asymmetry may therefore provide a non-invasive framework for investigating cortico-subcortical organisation in healthy and clinical populations.

