## Supplementary Materials for "Laterality of subcortical structures predicts spontaneous brain dynamics"

### Cluster-based permutation analysis

To identify spatially contiguous sensor clusters exhibiting significant associations between subcortical structural asymmetry and lateralised oscillatory power, we employed a non-parametric cluster-based permutation procedure implemented in Python using MNE-Python together with custom analysis code.

For each frequency band (Delta, Theta, Alpha and Beta), subcortical structure (thalamus, caudate, putamen, pallidum, hippocampus, amygdala and nucleus accumbens), and sensor type (magnetometers and combined planar gradiometers analysed separately), Spearman's rank correlations were computed between the lateralisation index of the subcortical volume and the lateralised power measured at each sensor pair. Because sensor-level inference involves multiple spatially correlated statistical tests, significance was assessed using cluster-based permutation testing.

The observed Spearman correlation coefficients were first transformed into *t*-statistics according to

$$t=\rho\sqrt{\frac{n-2}{1-\rho^{2}}}$$

where $\rho$ denotes the Spearman correlation coefficient and *n* is the number of participants included in the analysis. This transformation allowed a common threshold to be defined using the theoretical *t*-distribution. Sensors exceeding the two-tailed critical threshold (*α* = 0.05) were considered candidate significant sensors.

Spatial clusters were then formed using the sensor adjacency matrix generated from the MEG sensor layout. Sensor neighbourhoods were obtained using the MNE sensor adjacency functions, which define neighbouring sensors according to the physical arrangement of the MEG array. Only sensors exceeding the statistical threshold contributed to clustering. Connected components within the thresholded adjacency graph were identified using the *connected_components* algorithm from *scipy.sparse.csgraph*, thereby grouping neighbouring significant sensors into spatial clusters. Consequently, each cluster consisted of adjacent sensors whose transformed correlation values all exceeded the initial statistical threshold.

For each observed cluster, a cluster-level statistic was calculated as the sum of the *t*-statistics across all sensors belonging to that cluster,

$$T_{cluster}=\sum_{i=1}^{N_{c}} t_{i},$$

where $N_{c}$ is the number of sensors within the cluster. Positive and negative clusters were treated separately, allowing clusters reflecting positive and negative structure-function associations to be evaluated independently.

Statistical significance of the cluster-level statistics was determined using permutation testing. For each permutation, the correspondence between participants' subcortical lateralisation indices and their lateralised MEG power values was randomly shuffled while preserving the spatial structure of the MEG data. Sensor-wise Spearman correlations were recomputed for the shuffled data, transformed to *t*-statistics using the same equation, thresholded using the identical critical *t*-value, and spatial clusters were reconstructed using the same sensor adjacency matrix. Cluster statistics were calculated for each permutation as the sum of *t*-statistics within each cluster.

To control the family-wise error rate, only the largest positive cluster statistic and the most negative cluster statistic from each permutation were retained, generating empirical null distributions of maximum positive and minimum negative cluster masses. Following 1,000 permutations, significance thresholds were defined as the 95th percentile of the positive null distribution and the 5th percentile of the negative null distribution. An observed cluster was considered statistically significant if its cluster statistic exceeded the corresponding empirical threshold.

Only sensors belonging to significant clusters after permutation testing were retained for subsequent visualisation and spectral analyses.

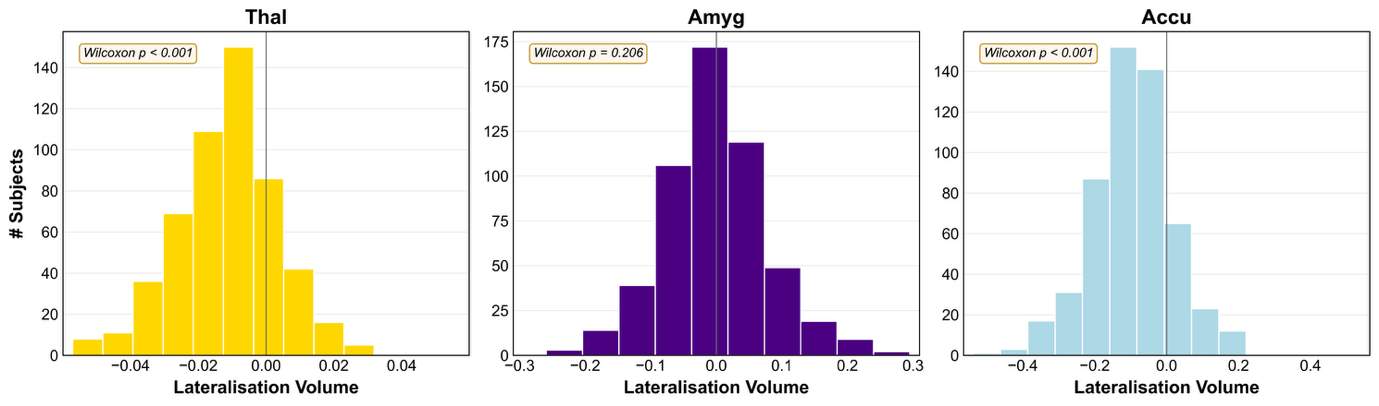

Supplementary figure 1. Distribution of Thalamus (A), Amygdala (B), and Nucleus Accumbenc (C) lateralised volume across participants (N = 532). Positive values indicate greater right than left volume, whereas negative values indicate greater left than right volume. Thalamus and Nucleus Accumbenc showed significant leftward lateralisation (Wilcoxon rank sum p < 0.001 for both structures).
